# Dysregulated Platelet Function in Patients with Post-Acute Sequelae of COVID-19

**DOI:** 10.1101/2023.06.18.545507

**Authors:** Anu Aggarwal, Tamanna K. Singh, Michael Pham, Matthew Godwin, Rui Chen, Thomas M. McIntyre, Alliefair Scalise, Mina K. Chung, Courtney Jennings, Mariya Ali, Hiijun Park, Kristin Englund, Alok A. Khorana, Lars G. Svensson, Samir Kapadia, Keith R. McCrae, Scott J. Cameron

## Abstract

**Background:** Post-acute sequelae of COVID-19 (PASC), also referred as Long-COVID, sometimes follows COVID-19, a disease caused by SARS-CoV-2. While SARS-CoV-2 is well-known to promote a prothrombotic state, less is known about the thrombosis risk in PASC.

**Aim:** Our objective was to evaluate the platelet function and thrombotic potential in patients following recovery from SARS-CoV-2 with clear symptoms of PASC.

**Methods:** PASC patients and matched healthy controls were enrolled in the study on average 15 months after documented SARS-CoV-2 infection. Platelet activation was evaluated by Light Transmission Aggregometry (LTA) and flow cytometry in response to platelet surface receptor agonists. Thrombosis in platelet-deplete plasma was evaluated by Factor Xa activity. A microfluidics system assessed thrombosis in whole blood under shear stress conditions.

**Results:** A mild increase in platelet aggregation in PASC patients through the thromboxane receptor was observed and platelet activation through the glycoprotein VI (GPVI) receptor was decreased in PASC patients compared to age- and sex-matched healthy controls. Thrombosis under shear conditions as well as Factor Xa activity were reduced in PASC patients. Plasma from PASC patients was an extremely potent activator of washed, healthy platelets – a phenomenon not observed when stimulating healthy platelets after incubation with plasma from healthy individuals.

**Conclusions:** PASC patients show dysregulated responses in platelets and coagulation in plasma, likely caused by a circulating molecule that promotes thrombosis. A hitherto undescribed protective response appears to exists in PASC patients to counterbalance ongoing thrombosis that is common to SARS-CoV-2 infection.

## Introduction

COVID-19, caused by the severe acute respiratory syndrome coronavirus 2 (SARS-CoV-2), may have residual effects following recovery. One third of infected patients are asymptomatic, while others experience pulmonary and cardiovascular symptoms leading to morbidity and mortality (1). The risk of both severe and persistent symptoms of SARS-CoV-2 is higher males, in the advanced-age patient population, in patients requiring immunosuppression, or with chronic kidney disease (2). The risk of severe and persistent symptoms of SARS-CoV-2 remains despite vaccination (3, 4).

Post-acute sequelae of COVID-19 (PASC) is an recognized multi-systemic condition characterized by persistent symptoms four weeks beyond the initial SARS-CoV- 2 infection. While acute respiratory distress syndrome (ARDS) is a feared consequence of severe COVID-19, many acutely ill patients are reported to have thromboembolic complications including deep vein thrombosis (DVT) and pulmonary embolism (PE). The incidence of venous thromboembolic (VTE) complications may be as high as 40% (5, 6).

Observational data suggests that PASC patients develop or have persistent symptoms long after recovery that were initially labeled Long COVID by the Centers for Disease Control (CDC) and the National Institutes of Health (NIH). The National Institute for Health and Care Excellence (NICE) now defines PASC as post-infection symptoms beyond 4-12 weeks (7-9). Patients blighted by the specter of PASC show structural and functional organ derangements involving the respiratory, cardiovascular, neurological, genitourinary, gastrointestinal, and musculoskeletal systems. PASC symptoms typically include fatigue, dyspnea, chest pain/tightness, headache, cough, difficulty concentrating, and altered sensorium. Poor memory and concentration are the most frequent symptoms reported, closely followed by cognitive impairment, sleep disturbance, and post-traumatic stress disorder (PTSD) (9-13).

Elevated pro-thrombotic markers including D-dimer, C-reactive protein, and neutrophil extracellular trap (NET) formation have been detected in acute SARS-CoV-2 infection. In addition, many PASC patients complain of dyspnea on exertion, which leads to physical inactivity and may further precipitate thromboembolic complications. Virus-induced endothelial injury of severe COVID-19 patients initiates a state of dysregulated coagulation. Increased blood von Willebrand Factor (VWF), likely from both endothelial cell and platelet activation, promotes hypercoagulability and thrombosis in patients with COVID-19 (7, 8, 14).

Platelets have an important hemostatic function to prevent excessive bleeding (hemostasis) alongside an important role in the host immune response to invading viral and microbial infections, especially SARS-CoV-2 (15). In a large observational study, no additional risk for VTE after hospitalization with acute SARS-CoV-2 infection was noted (16). Contrasting this, the MICHELLE study revealed a protective effect of an oral Factor Xa inhibitor on discharged patients by protecting them against DVT if they were previously hospitalized, even without an initial thrombotic event (17). These findings raise the possibility that following SARS-CoV-2 recovery, patients may have ongoing functional impairment in the coagulation cascade or abnormal platelet function. The Aim of this study was to evaluate both platelet function and thrombotic potential in whole blood, in platelet-deplete plasma, and in isolated platelets in patients following SARS-CoV-2 but with clear PASC symptoms.

## Methods

### Human Subjects

26 patients with Post-Acute Sequelae of SARS-CoV-2 infection (PASC) were enrolled based on a confirmed Covid-19 diagnosis and with persistent symptoms, including shortness of breath, fatigue, brain fog, dyspnea on exertion, mood disturbance, weakness, and memory impairment. PASC patients were compared to 19 matched healthy controls. Patients and healthy subjects were recruited only after informed consent by a research coordinator not involved in clinical care. All study procedures were conducted in accordance with the Declaration of Helsinki and approved by a local Institutional Review Board (IRB) protocol. Human platelet physiological studies must be conducted within 90 minutes in order for meaningful data to be obtained. These studies are labor-intensive and, in their entirety, cannot be conducted simultaneously for every patient within 90 minutes. The volume of blood required to conduct studies in whole blood, in platelet-rich plasma (PRP), in washed platelets, and in platelet deplete plasma is also substantial and exceeds the blood volume permitted to be drawn in the IRB protocol. Therefore, it is impossible to conduct every experiment in every enrolled PASC patient or healthy individual. Rather, the studies were conducted in a manner that was practically required as patients as PASC were identified.

### Cross-activation study using Flow-cytometry

To check whether plasma from PASC patients would modify platelet activation from healthy donors, we performed cross-activation tests. Platelet depleted plasma (20μl) from the PACS patients (n=18) as well as age- and sex-matched healthy controls (n=20) was added to the isolated washed platelets (80μl) from healthy controls (each performed in triplicate) and incubated for 20 minutes at room temperature. Platelets were stimulated with agonists: Thrombin Receptor Activator Peptide 6 (TRAP6; 5μM), Thromboxane A2 (TXA2) analogue U46619 (1μM), Adenosine diphosphate (ADP; 0.1μM), or Collagen Related Peptide (CRP; 0.1μg/mL) for 15 minutes. Samples were next stained with phycoerythrin (PE)-conjugated anti-CD62P (p-selectin) for 30 minutes at room temperature in darkness to prevent fluorophore bleaching before fixing with 4% paraformaldehyde. Fixed samples were read by flow cytometry (BD Accuri C6 Plus) to acquire 20000 events and data was analyzed using FlowJo software.

### Antibody Profiling

Antibody profiling was performed by CDI labs using the HuProt 4.0 array. Briefly, sequence-confirmed plasmids, which are used to make GST-purified recombinant proteins in yeast. After purification, GST-tagged proteins are piezoelectrically printed on glass slides in duplicate, along with control proteins (GST, BSA, Histones, IgG, etc.). The slides are barcoded for tracking and archiving. Each microarray batch is routinely evaluated by anti-GST staining to demonstrate quality of expression and printing. The new HuProt v4.0 consists of unique human proteins, isoform variants, and protein fragments of the 19,613 canonical human proteins described in the Human Protein Atlas (18-21). We focused on protein exclusively expressed in platelets, highly expressed in platelets, blood proteins known to be involved in the alterations in coagulation as noted. Content includes major functional classes such as intracellular proteins, membrane proteins, enzymes, secreted proteins, transcription factors, transporters, GPCRs, cytokines, immune receptors, immune checkpoints, CD markers, ion channels, cytosolic proteins, nuclear receptors. To measure antibody binding, slides are overlaid with a 1:100 dilution of serum, and stained with anti-IgG (red) and anti-IgM (green) secondary antibodies. A heat map was generated to show each antibody concentration expressed as standard deviations (SD) above or below the mean concentration found in healthy individuals.

### Light Transmission Aggregometry

**(LTA)** Blood from healthy controls and PASC patients was collected in sodium-citrate tubes after obtaining informed consent. Within 30 minutes to 1 hour of collection, anticoagulated whole blood was centrifuged at 200g for 15 minutes at room temperature (RT) to obtain platelet rich plasma (PRP). PRP was collected in 15mL falcon tubes and Platelet poor plasma (PPP) was obtained from the remaining sample by re-centrifugation at 2500 g for 10 minutes. PPP was used to initiate baseline optical density. Platelet Aggregation was tested following stimulation with agonists: ADP (1μM), TRAP-6 (5μM), U46619 (5μM) and CRP (1μg/mL). Optical density changes were recorded after the addition of the agonists on Aggregometer (CHRONO-LOG 700) for 6 minutes while stirring the sample at 1200 rpm at 37°C.

### Alpha granule exocytosis with Flow-cytometry

Platelets were isolated from whole blood collected from healthy controls and PASC patients in sodium-citrate tubes. Briefly, whole blood was centrifuged at 200g for 15 minutes at RT and then PRP was collected in the 15mL falcon tubes. Collected PRP was centrifuged again at 200g for 10 minutes to remove remaining WBCs and RBCs. Platelets were washed using Tyrode’s buffer in the 1:1 ratio with the addition of Prostaglandin I2 (PGI2; 10nM). Washed platelets were re-suspended in 10mL of Tyrode’s buffer. Washed platelets from PASC patients and healthy controls were stimulated using different concentrations of agonist: TRAP-6 (1, 5, 10 and 20μM), U46619 (0.1, 0.5, 1.0, 10μM), ADP (0.01, 0.1, 1.0, 10μM) and CRP (0.05, 0.1, 0.5, 1.0μg/mL) for 15 minutes and then incubated with phycoerythrin (PE)-conjugated anti-CD62P for 30 minutes at RT. Samples were fixed with 4% paraformaldehyde. P-Selectin expressed on the surface on the platelets signifies an activated state. Samples were read by flow cytometry (BD Accuri C6 Plus) to acquire 20000 events. Data were analyzed using FloJo software.

### Total Thrombus formation Analysis System (T-TAS01)

The T-TAS01 microfluidic chamber system was used and the results were compared to healthy controls. T-TAS01 is a microchip flow-chamber system that measures thrombus formation under blood flow conditions over time. Whole blood collected in the sodium-citrate tube was used for the test. Briefly, before measurement 20μL of the CaCTI reagent (Corn trypsin inhibitor) was added to 480μL of whole blood and then this blood was mixed gently with the pipette. Then 450μL of mixed blood was loaded into the reservoir of the AR chip reservoir coated with both collagen and thromboplastin (tissue factor) with a shear rate of 600 s^-1^. As coagulation and platelet activation progress, thrombosis is noted as pressure rises in the channel following shear rate-mediated activation of blood in the microchip channel. These pressure changes are monitored by a pressure sensor upstream of the chamber and are defined as The Occlusion Start Time (OST), and the Occlusion Time (OT). The difference between OT and OST was calculated for each subject. Furthermore, to check the pro-thrombotic activity of the plasma of the PASC patients, 200μL plasma from the PASC patients and healthy controls was incubated with 800μL whole blood from healthy control for 30 minutes. Then this spiked whole blood with plasma was loaded into the reservoir of the AR chip.

### Plasma Factor Xa activity

Factor Xa (FXa) is the activated form of the coagulation factor X. Factor X, a serine endopeptidase plays an important role in the coagulation pathway. It converts prothrombin into active thrombin by complexing with activated co-factor V in the prothrombinase complex. The Factor Xa Activity assay kit from Abcam (ab204711) was used to measure the Factor Xa activity in the plasma of the PASC patients compared to age- and sex-matched healthy controls. The assay was performed according to the manufacturer’s recommendations.

### Statistical Analysis

All data were interrogated for normalcy using the Shapiro-Wilks test. Normally distributed continuous variable group differences were assessed by the student 2-tailed Student’s t-test and skewed data assessed by the Mann-Whitney *U* test. Analyses were conducted using GraphPad Prism 7 (GraphPad Software). For Gaussian-distributed data in 3 or more groups, 1-way ANOVA then the Bonferroni multiple comparisons test was used, otherwise the Kruskal–Wallis test followed by Dunn post-test was used. Significance was accepted as a *P* value of less than 0.05.

## Results

### Baseline Study Population

26 patients with PASC (mean age 49.9) were recruited from the pulmonary medicine clinic in Northeast Ohio and compared with 19 healthy volunteers (mean age 41.4). Healthy recruits or PASC patients were comparable, showing no significant difference between groups with respect to sex, BMI, and medical co-morbidities of PASC patients who were more commonly smokers (**Table 1**). Patients with PASC presented on average 501 days ± 69 days after the last detection of a SARS-CoV- 2 amplicon by polymerase chain reaction (PCR). Of the 26 common PASC symptoms in our patient population, most described fatigue (76.9%), brain fog (76.9%), shortness of breath at rest (69.2%), and exertional dyspnea (69.2%) (**Supplemental Table 1**).

**Table 1.** Demographics for PASC Patients and Healthy Controls.

| Demographics | Healthy (n=19) |  | PASC (n=26) |  | P-Value |
| --- | --- | --- | --- | --- | --- |
|  | N | % | N | % |  |
| **<u>Characteristics</u>** |  |  |  |  |  |
| Mean Age (± SEM) | 41.4 ± 2.89 | | 49.9 ± 3.61 | | 0.090 |
| 18-27 years | 4 | 21.0% | 3 | 11.5% | 0.30 |
| 28-40 years | 4 | 21.0 % | 7 | 26.9 % | 0.76 |
| 41-60 years | 9 | 47.3 % | 5 | 19.2 % | 0.51 |
| 61-82 years | 2 | 10.5 % | 11 | 42.3 % | 0.62 |
| Male | 4 | 21.0 % | 7 | 35.0 % | 0.73 |
| Female | 15 | 78.9 % | 19 | 73.0 % |  |
| BMI (± SEM) | 29.5 ± 1.29 | | 30.9 ± 2.17 | | 0.61 |
| Smoker | 1 | 7.7 % | 13 | 50.0 % | 0.002 |
| **<u>Medical History</u>** |  |  |  |  |  |
| Hypertension | 4 | 21.0 % | 6 | 23.0 % | 0.99 |
| Hyperlipidemia | 2 | 10.5 % | 5 | 19.2 % | 0.68 |
| History of CVD | 1 | 5.2 % | 3 | 11.5 % | 0.62 |
| Diabetes | 2 | 10.5 % | 1 | 3.8 % | 0.56 |
| History of Pulmonary Embolism | 0 | 0.0 % | 1 | 3.8 % | 0.99 |
| History of Thrombosis | 1 | 5.2 % | 0 | 0.0 % | 0.42 |
| **<u>Medications</u>** |  |  |  |  |  |
| Platelet inhibitors | 1 | 5.2 % | 6 | 23.0 % | 0.21 |
| Aspirin | 1 | 5.2 % | 5 | 19.2% |  |
| DAPT (Dual Antiplatelet Therapy) |  |  | 1 | 3.8% |  |
| Anticoagulants | 0 | 0.0 % | 2 | 7.6 % | 0.50 |
| Warfarin |  |  | 1 | 3.8% |  |
| Apixaban |  |  | 1 | 3.8% |  |
| Statins | 2 | 15.4 % | 5 | 19.2 % | 0.68 |
| NSAIDs | 6 | 31.5 % | 4 | 15.38 % | 0.28 |
| BP Medications | 4 | 21.0 % | 1 | 3.8 % | 0.14 |

### Platelet Reactivity in PASC patients

When platelet-deplete plasma from healthy individuals or PASC patients was added to healthy, twice washed platelets, there was a surprisingly profound activation of platelets following stimulation with all platelet receptor agonists tested caused by PASC patient plasma incubation (**Fig. 1**). Given that immunothrombosis is common in COVID-19 and the complement system likely contributes to these responses (22-24), we heated and inactivated plasma from PASC donors at 55°C and repeated the experiments. We observed that platelets were persistently activated if spiked with room temperature plasma or heat-inactivated plasma prior to agonist stimulation from PASC patients suggesting that complement activation is unlikely to be a driver in the augmented platelet response observed (**Supplemental Fig. S1**).

**Fig. 1.**
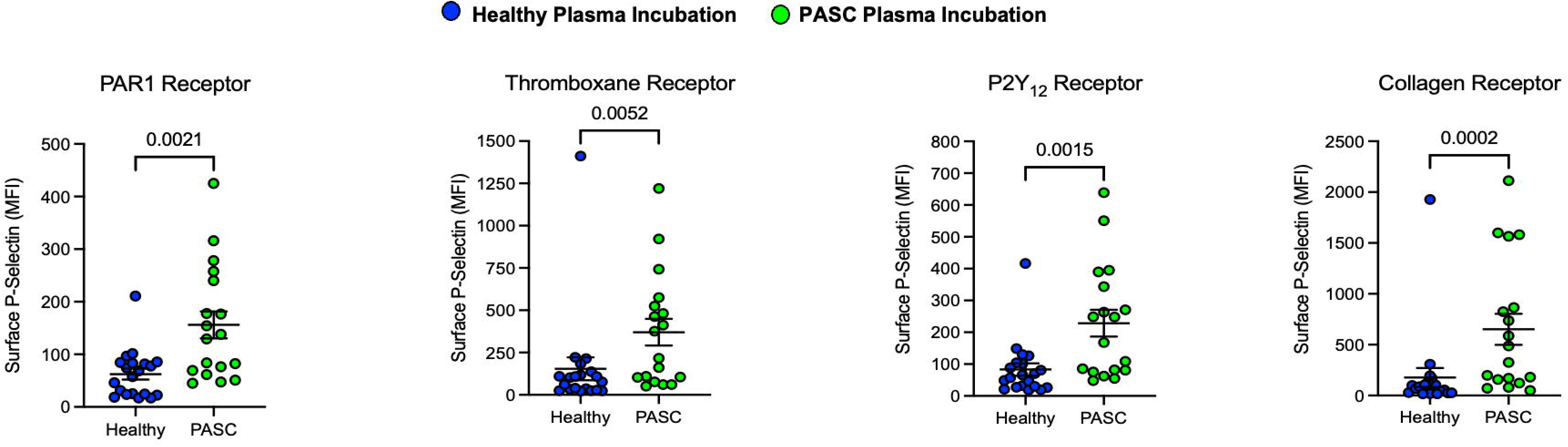
Activation of Platelets by PASC but not Healthy Plasma. Isolated, washed platelets were pre-incubated for 30 minutes with 20 µL of age- and sex-matched healthy plasma or PASC plasma, then stimulated with TRAP-6, 5 µM (PAR1), U46619, 1 µM (Thromboxane receptor), ADP, 0.1 µM (P2Y12 Receptor), or Collagen-Related Peptide, CRP, 0.1 µg/mL (GPVI Receptor) for 15 minutes. Platelet reactivity was assessed by FACS by surface CD62P (P-selectin) using a tagged antibody. Data are represented as mean ± SEM. Group differences were assessed by the Mann-Whitney U test, n=20 plasma isolated for healthy individuals and n=18 plasma isolates for PASC patients. Washed platelets from a healthy 25 year-old male.

Platelet aggregation in PRP was evaluated by LTA after stimulating platelets with surface receptor agonists for the P2Y12 receptor (ADP), the thromboxane receptor (U46619), the GPVI receptor (CRP), and PAR1 (TRAP-6). Overall, platelet aggregation by LTA in PASC was similar to healthy individuals except through the thromboxane receptor which demonstrated increased reactivity following stimulation with the agonist U46619 (87%±3 for PASC vs. 72%±4 for healthy subjects, p=0.03) (**Fig. 2A-D**). To distinguish platelet function that may be altered by mediators in PRP, these studies were repeated in a dose-response manner after isolating and washing platelets twice, then assessing platelet reactivity by alpha granule exocytosis by appearance of P-selectin on the surface of the platelet as reported previously (25, 26). When washed platelets from healthy volunteers and PASC patients were stimulated with platelet surface receptor agonists, we found similar reactivity to healthy individuals through the following receptors: P2Y12, thromboxane, PAR 1 (**Fig. 3A-C**). Curiously, platelet reactivity through GPVI was markedly decreased in PASC patients compared with healthy controls (**Fig. 3D**).

**Fig. 2A.**
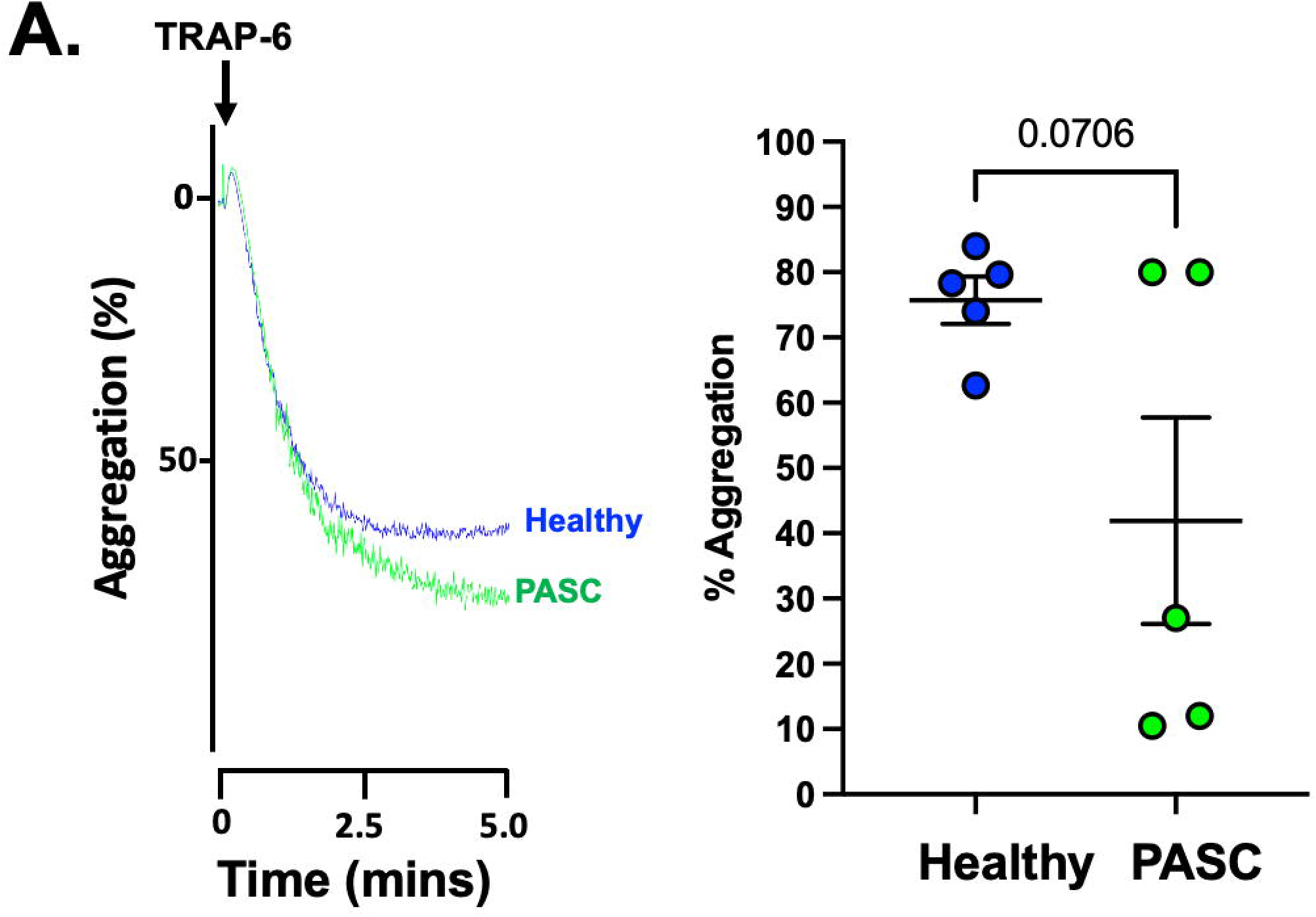
Platelet Activation in PASC by Aggregometry. Blood was drawn from healthy individual or PASC patients and PRP was isolated prior to stimulation with TRAP-6, 5µM until platelet aggregation was noted. Platelet aggregation was evaluated by LTA and data reported as mean % maximum aggregation ± SEM. A representative aggregometry tracing is shown. Group differences were assessed by the Mann Whitney *U* test, n=5 for healthy individuals and n=5 for PASC.

**Fig. 2B.**
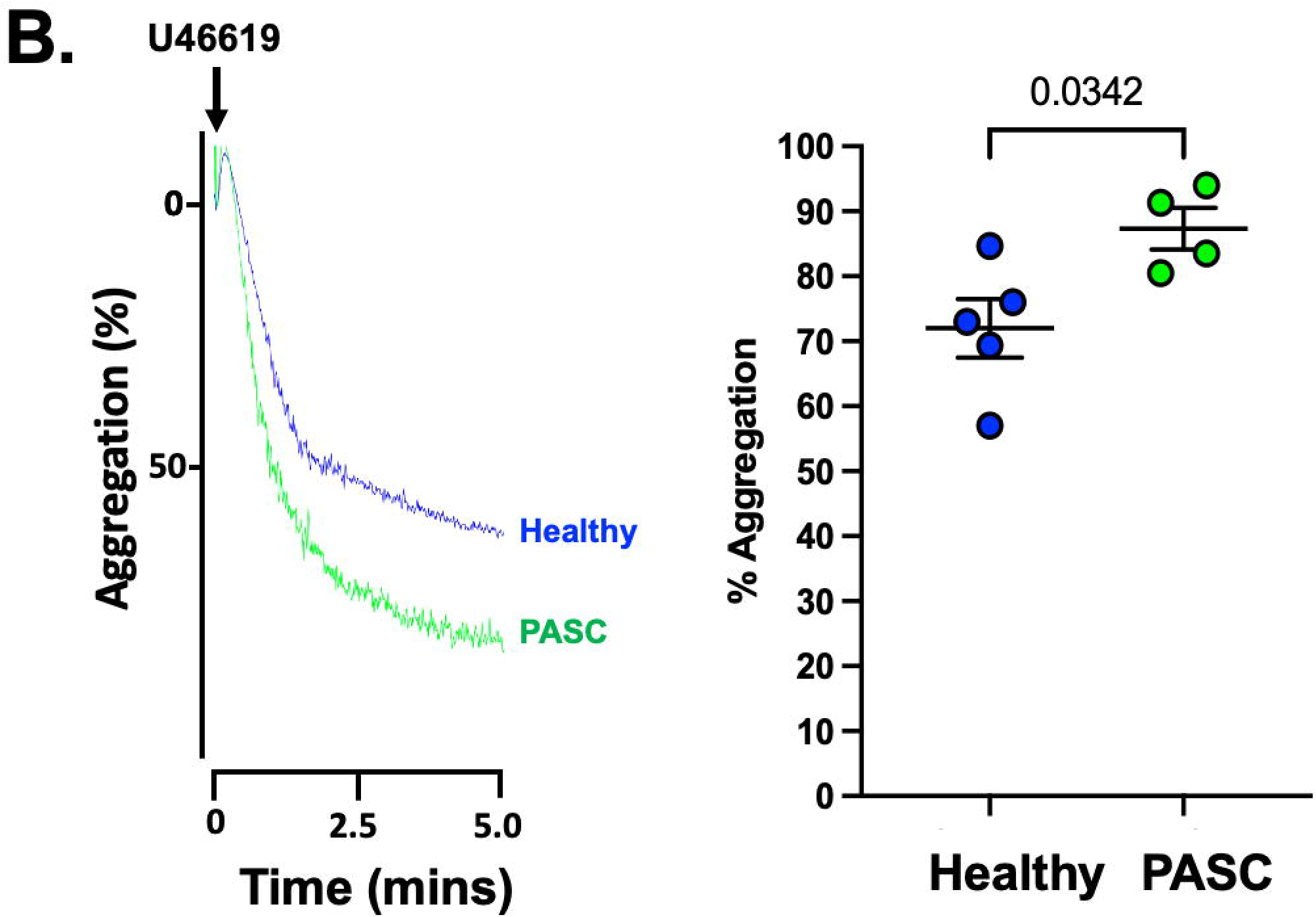
Platelet Activation in PASC by Aggregometry. Blood was isolated from healthy individual or PASC patients and PRP was isolated prior to stimulation with U466119, 5µM until platelet aggregation was noted. Platelet aggregation was evaluated by LTA and data reported as mean % maximum aggregation ± SEM. A representative aggregometry tracing is shown. Group differences were assessed by the student’s t-test, n=5 for healthy individuals and n=5 for PASC.

**Fig. 2C.**
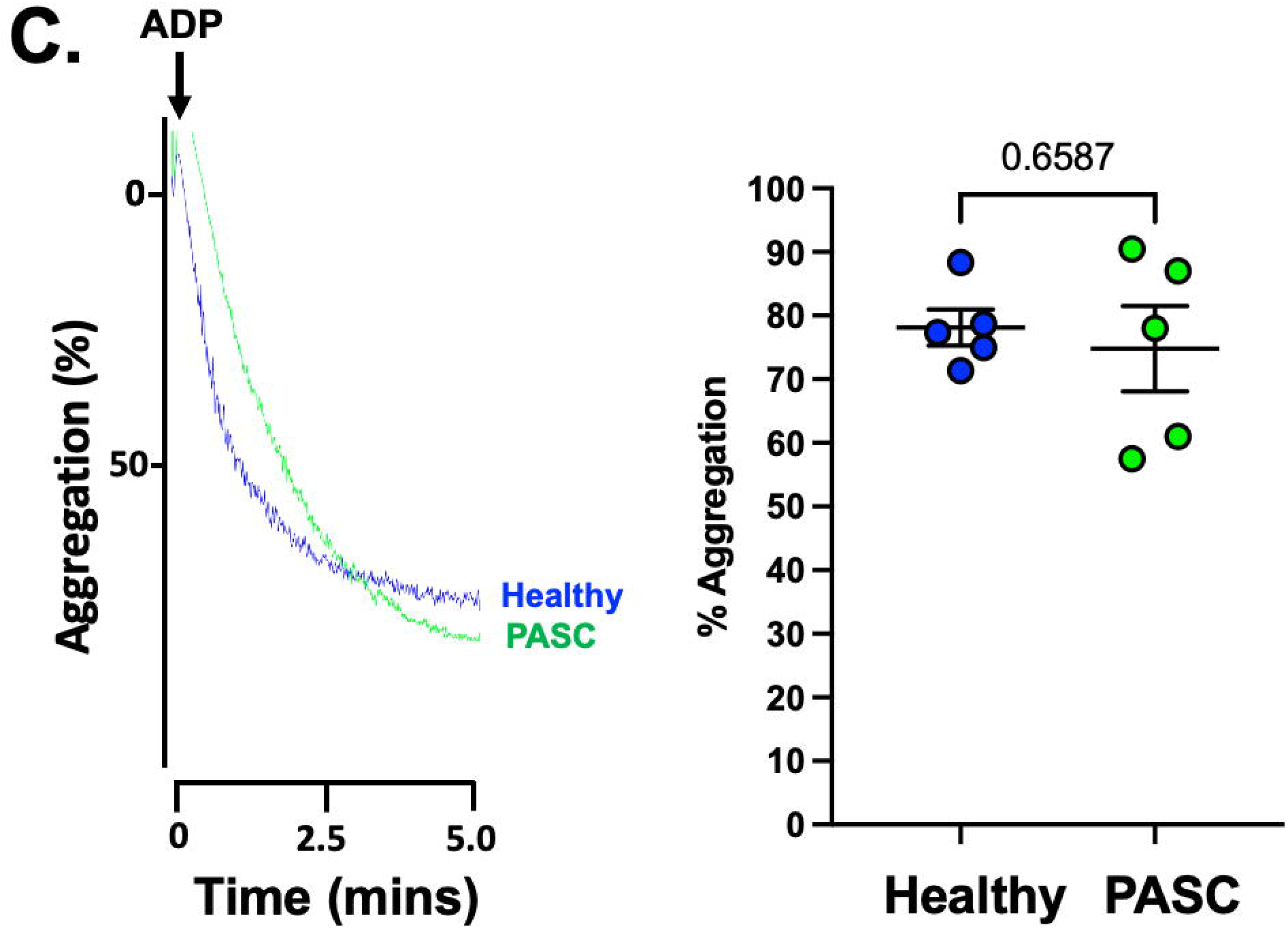
Platelet Activation in PASC by Aggregometry. Blood was isolated from healthy individual or PASC patients and PRP was isolated prior to stimulation with ADP, 1 µM until platelet aggregation was noted. Platelet aggregation was evaluated by LTA and data reported as mean % maximum aggregation ± SEM. A representative aggregometry tracing is shown. Group differences were assessed by the Mann Whitney *U* test, n=5 for healthy individuals and n=5 for PASC.

**Fig. 2D.**
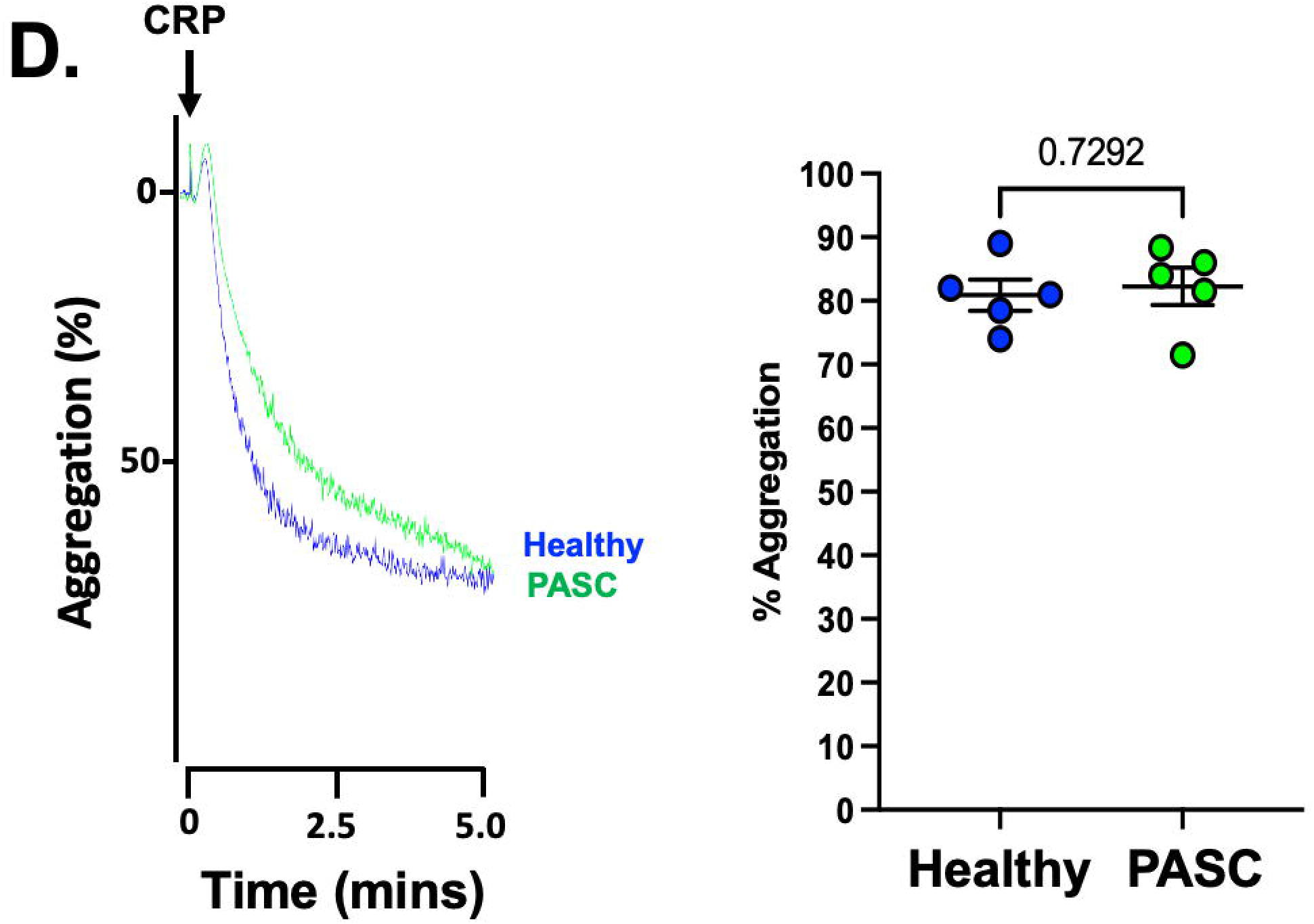
Platelet Activation in PASC by Aggregometry. Blood was isolated from healthy individual or PASC patients and PRP was isolated prior to stimulation with CRP, 1 µg/mL until platelet aggregation was noted. Platelet aggregation was evaluated by LTA and data reported as mean % maximum aggregation ± SEM. A representative aggregometry tracing is shown. Group differences were assessed by the Mann Whitney *U* test, n=5 for healthy individuals and n=5 for PASC.

**Fig. 3A.**
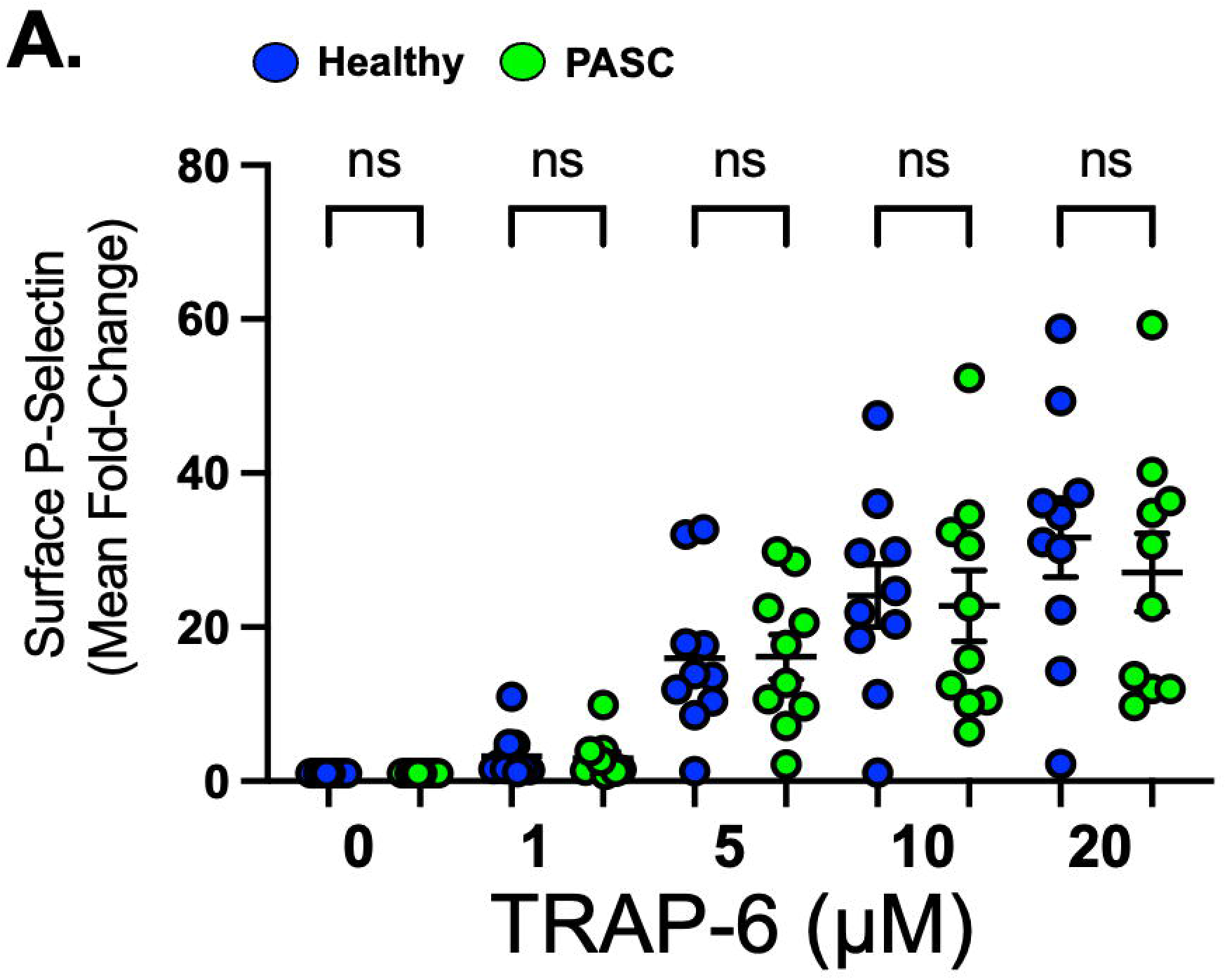
Platelet Activation in PASC by Flow Cytometry. Isolated, washed platelets were stimulated with TRAP-6 for 15 minutes, then reactivity was assessed by FACS by fold-change in surface CD62P (P-Selectin) using a tagged antibody. Data are represented as mean fold-change ± SEM. Group differences were assessed by the Kruskal-Wallis test, n=10 for healthy individuals and n=9 for PASC. NS=not significant.

**Fig. 3B.**
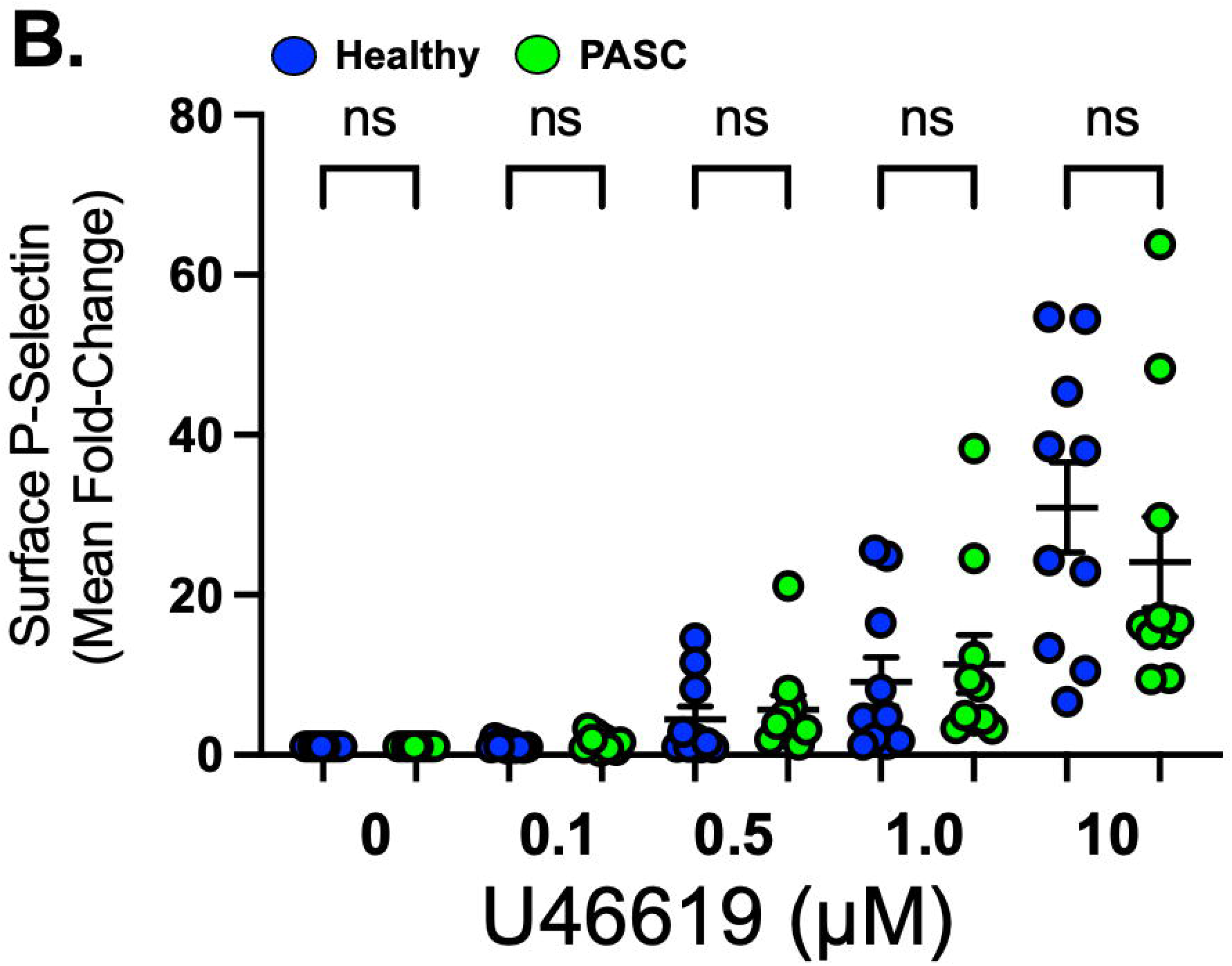
Platelet Activation in PASC by Flow Cytometry. Isolated, washed platelets were stimulated with U46619 for 15 minutes, then reactivity was assessed by FACS by fold-change in surface CD62P (P-selectin) using a tagged antibody. Data are represented as mean fold-change ± SEM. Group differences were assessed by ANOVA, followed by the Bonferroni correction, n=10 for healthy individuals and n=10 for PASC.

**Fig. 3C.**
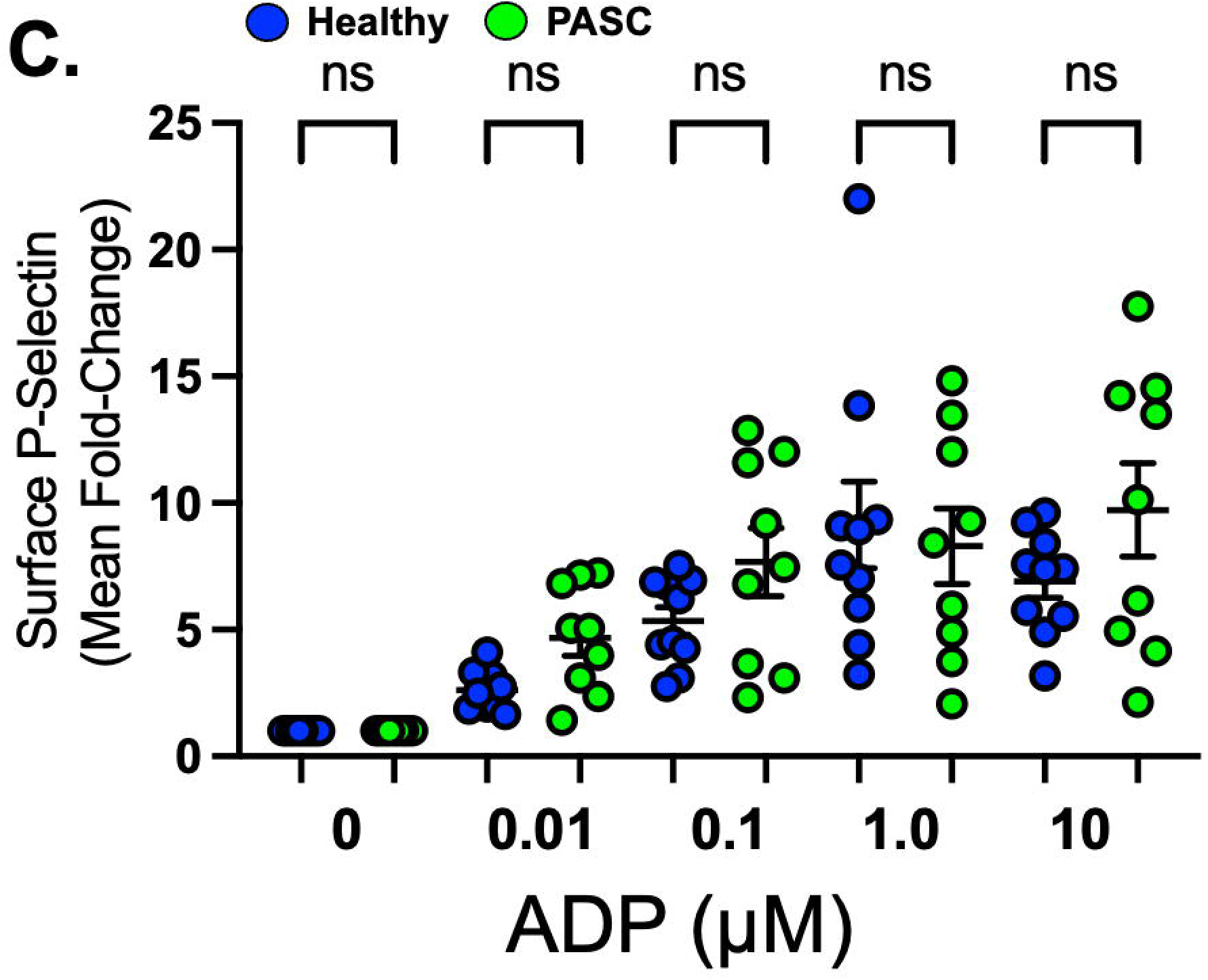
Platelet Activation in PASC by Flow Cytometry. Isolated, washed platelets were stimulated with ADP for 15 minutes, then reactivity was assessed by FACS by fold-change in surface CD62P (P-selectin) using a tagged antibody. Data are represented as mean fold-change ± SEM. Group differences were assessed by ANOVA, followed by the Bonferroni correction, n=10 for healthy individuals and n=9 for PASC.

**Fig. 3D.**
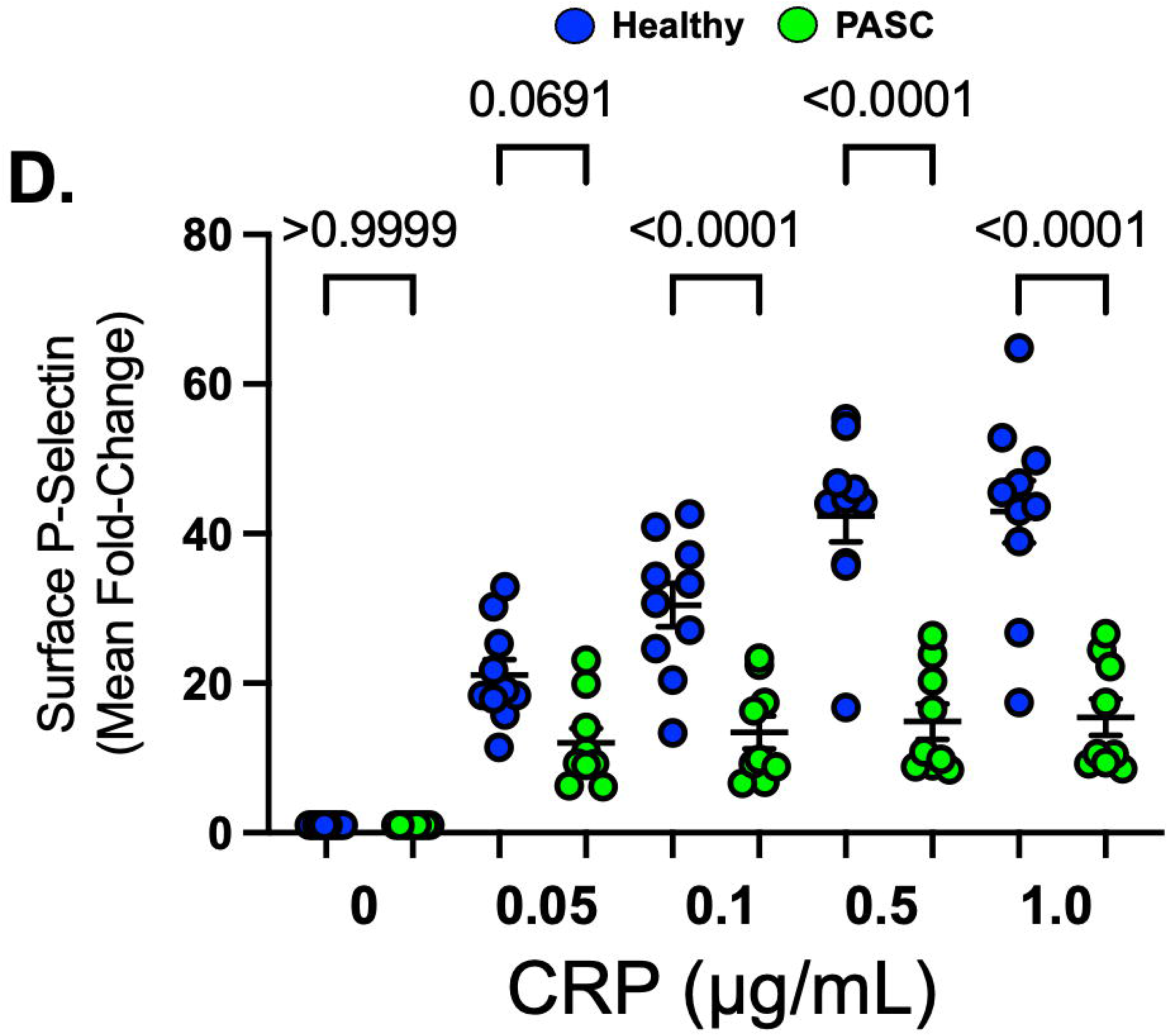
Platelet Activation in PASC by Flow Cytometry. Isolated, washed platelets were stimulated with collagen-related peptide (CRP) for 15 minutes, then reactivity was assessed by FACS by fold-change in surface CD62P (P-selectin) using a tagged antibody. Data are represented as mean fold-change ± SEM. Group differences were assessed by ANOVA, followed by the Bonferroni correction, n=10 for healthy individuals and n=9 for PASC.

To test the hypothesis that changes in platelet function as observed by alpha granule exocytosis augmented in PASC following platelet thromboxane receptor stimulation, **Fig. 2B**) or platelet aggregation (decreased in PASC following platelet GPVI stimulation, **Fig. 3D**) are a consequence of changes in platelet surface receptor expression, we performed flow cytometry using receptors tagged with a fluorophore, ultimately finding no difference (**Supplemental Fig. S2**).

### Whole blood and platelet-independent coagulation in PASC patients

Whole blood was drawn from patients who were healthy or with PASC, then passed through a 80 μm channel in a microfluidics system at 600 dynes (s^-1^) shear stress. This assay was specifically chosen because thrombosis in most patients with SARS-CoV-2 occurs in the venous vasculature, and the assay utilizes tissue factor and collagen to initiate thrombosis, which simulates endothelial damage common to SARS-CoV-2. Overall, the time to pressure increase in the microchannel (kPa), which is a signature of *ex vivo* blood clotting as coagulated blood diminishes flow, was prolonged in PASC patients compared with healthy conditions (occlusion time-occlusion start time, 94±8 vs. 73±8 seconds, respectively, P=0.02) (**Fig. 4 A-B**). To test whether a mediator in plasma is responsible for the difference in coagulation observed in PASC, whole blood from a healthy individual was spiked with platelet-deplete plasma from a PASC patient was compared to spiking platelet-deplete plasma from a healthy individual. This showed no difference between occlusion time and occlusion start time (62±4 healthy plasma vs. 66±8 PASC plasma, n=5 in each group, P=0.74). We next assessed Factor Xa activity in platelet-deplete plasma which represents the point of convergence between the extrinsic and the intrinsic arms of the coagulation cascade. We observed Factor Xa activity was reduced in PASC vs. healthy individuals (23.6±5 vs. 50±8 ng/100µL, respectively; P=0.037) (**Fig. 4C**). While complement, which can increase platelet reactivity, is well-known to be denatured at 55°C, antibodies targeted against platelet proteins or the coagulation cascade may retain some function (27-29). We therefore performed a limited antibody array using serum from patients with PASC comparing this to healthy individuals for antibodies targeted against platelet proteins that could alter platelet function of antibodies targeted against circulating Factor II, Factor II Receptor, or Factor X. We did not find a significant difference in either IgM or IgG against those protein targets comparing PASC patients to healthy controls (**Supplemental Fig. S3**).

**Fig. 4.**
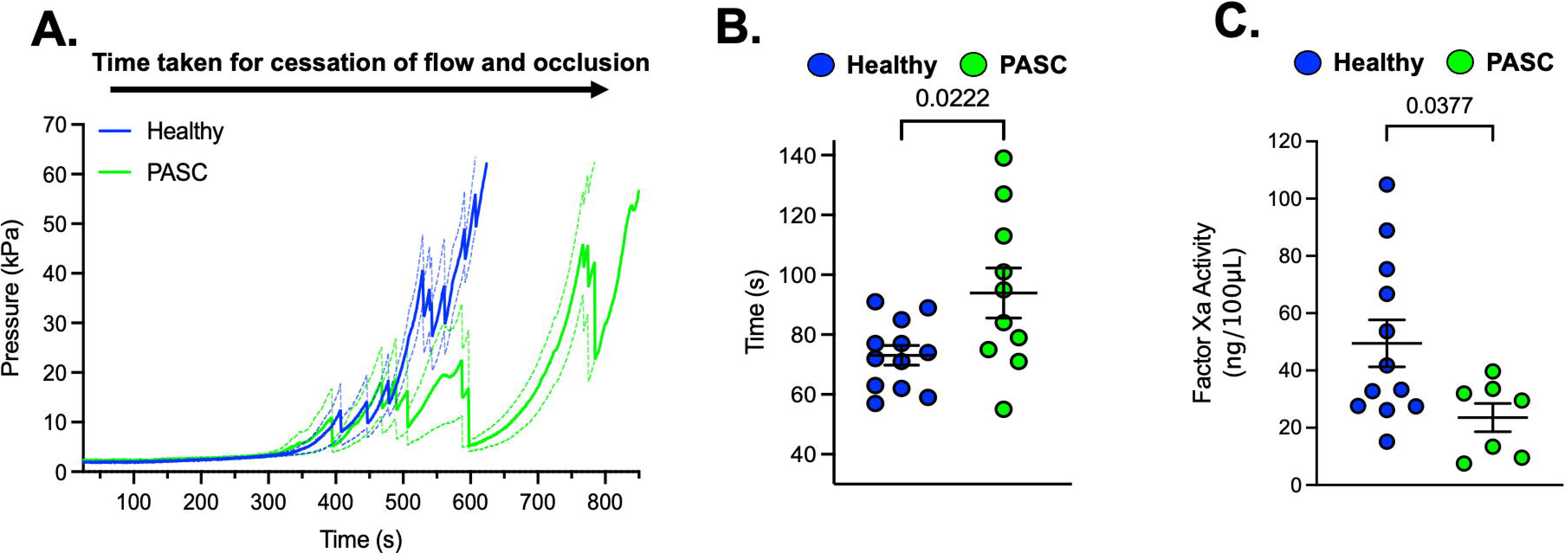
Whole Blood and Plasma Coagulation in PASC by Microfluidics and Factor Xa Activity. **A.**Blood was isolated and passed through a microfluidics chip using the Total Thrombus formation Analysis System (T-TAS01) at roughly venous shear stress (600 s-1). With clot generation, pressure in the microfluidics chamber (kPa) gradually increases until cessation of blood flow. Data are represented as mean ± SEM. **B.** The time from the start of occlusion to cessation of blood was calculated in seconds as mean ± SEM. Group differences were determined by the student’s t-test for PASC patients (n=10) or or healthy volunteers (n=12). **C.** Citrate plasma devoid of platelets was evaluated for Factor Xa activity using a chromogenic substrate in PASC patients (n=7) or healthy volunteers (n=10). Data are represented as mean ± SEM. Group differences were assessed by the student’s t-test, n=12 for healthy individuals and n=10 for PASC.

These experiments suggest there is a change in PASC plasma with dysregulated coagulation compared with healthy conditions that does not involve complement or circulating antibodies. Overall, the observation that washed platelets from PASC patients are relatively unchanged or less reactive than healthy platelets, and that platelet-deplete plasma in PASC is less likely to clot, suggests a potential adaptive response in the host long after COVID-19 infection, possibly to prevent thrombosis that is common in patients with acute COVID-19 infection.

## Discussion

SARS-CoV-2 is well-known to precipitate thrombotic events, especially DVT and PE (30, 31). Acute SARS-CoV-2 infection has also been shown to coincide with activated platelets by multiple independent groups (32-35). Our primary objective was to assess platelet function and thrombosis in patients with PASC given that rheologic manifestations of this syndrome are an under-developed area of investigation. Our major finding in patients more than one year following acute SARS-CoV-2 and with clear symptoms of PASC is dysregulated platelet function, with slight platelet activation through the thromboxane receptor, and attenuated activation through the GPVI receptor. We did not observe a change in platelet receptor signaling through PAR1 or P2Y12 in PASC using two well-validated techniques to interrogate platelet reactivity.

This study may be the first to report a functional defect in the coagulation cascade given our finding of reduced activated Factor Xa in platelet-deplete plasma in patients with PASC (33). Curiously, using radioligand binding technology or a similar chromogenic substrate for Factor Xa activation as we employ, others have reported that platelets and platelet-derived microparticles may be a source of Factor Xa activity in the circulation (36- 38). Moreover, a putative signaling mechanism between the platelet GPVI receptor and Factor Xa was previously reported (39). In addition, the profoundly under-reactive GPVI receptor identified by flow cytometry for alpha granule exocytosis using CRP as the agonist in our study complements the observation of prolonged time to thrombosis in human blood *ex vivo* under conditions of venous shear stress since collagen is present in the microfluidics channel which activates the GPVI receptor.

The most significant finding in our study is that platelet-deplete plasma in PASC patients ‘primes’ and markedly activates healthy, washed platelets through every receptor signaling pathway evaluated. We did not see this when platelet-deplete plasma from healthy controls was incubated with washed platelets before agonist stimulation. The hyper-reactive platelet effect caused by PASC plasma persisted even after heating plasma to 55°C suggesting the complement cascade or circulating antibodies are likely not responsible for augmented platelet activation. Using an antibody array, we could not find antibodies directed against platelet protein targets of coagulation proteins in the blood that could explain the observation in patients with PASC. Therefore, we offer the suggestion that a putative pro-thrombotic factor exists in the blood of patients with PASC, and its presence leads to an adaptive response in the host in which platelet reactivity and thrombosis are paradoxically tipped in the opposite direction, presumably as a protective response from thrombosis.

Our study in PASC patients differs from a recent letter by Martins-Gonçalves *et al.* in patients reporting at least one persistent respiratory symptom after acute hospitalization for COVID-19 and with persistent platelet hyperreactivity (40). Important differences in the enrolled subjects were noted. Firstly, while the average age of patients enrolled by Martins-Gonçalves *et al.* was similar to our cohort, the represented majority were men while 73% of our patients and 79% of our healthy controls were women. Platelet reactivity has been shown to differ between men and women in health and in cardiovascular disorders (26, 36, 41). Secondly, the control group employed by Martins-Gonçalves *et al*. tested negative for SARS-CoV-2 weekly for almost two years prior to enrollment, while several of our control comparators reported SARS-CoV-2 infection in the preceding two years but without residual symptoms at the time of enrollment and this was the reason why they were selected as controls. Lastly, Martins-Gonçalves evaluated platelet reactivity more in the convalescent setting 1-4 months after acute SARS-CoV-2 infection in 2021 whereas our PASC patients on average were evaluated >16 months after the last documented acute SARS-CoV-2 infection in 2023. Clinical outcomes and symptoms following SARS-Cov-2 infection appear to vary depending on vaccination status and the viral strain of the infected individual (42). This means that interpretation of data gained from patients evaluated during different waves of the COVID-19 pandemic require careful attention. Given that SARS-CoV-2 was demonstrated to induce morphological changes in platelets and promote apoptosis after viral internalization in an ACE2-dendent and ACE2-independent receptor internalization pathway (15), platelet reprogramming in patients with PASC may occur.

## Conclusion

In summary, patients who have long recovered from acute SARS-CoV-2 but with persistent symptoms under the constellation of PASC show evidence of dysregulated platelet reactivity, alterations in normal coagulation of platelet-deplete plasma, and diminished clotting in whole blood *ex vivo* under shear stress conditions. These events are likely a consequence of adapting to a platelet-activating factor in plasma of patients with PASC. The stimulus behind the exhausted platelet phenomenon, particularly through GPVI, in PASC patients, should be investigated and identification of the putative pro-thrombotic mediator should be prioritized given the implications for patients with PASC being considered as donors of blood and plasma products in future.

## Acknowledgements and Sources of Funding

We are grateful for funding from NHLBI HL128856 to SJC and HL158669 to TMM. Dr. Khorana acknowledges research support from the Sondra and Stephen Hardis Chair in Oncology Research. The research was, in part, funded by the National Institutes of Health (NIH) Agreement 1OT2HL156812 through the National Heart, Lung, and Blood Institute (NHLBI) CONNECTS program. The views and conclusions contained in this document are those of the authors and should not be interpreted as representing the official policies, either expressed or implied, of the NIH.

## Tables and Figures

**Table 1. Baseline Characteristics.** Patients with PASC (n=26, occurring at a mean duration of 501 ± 69 days after a diagnosis of COVID-19) were recruited for the study and compared with healthy volunteers (n=19). Data are reported as mean ± SEM. Differences between groups was calculated by the student’s t-test for continuous variables or Chi-Squared for dichotomous variables. BMI=Body Mass Index. CVD=Cardiovascular Disease. BP=Blood Pressure. NSAID=non-steroidal anti-inflammatory medication.

## Supplementary Tables and Figures

**Table S1. Persistent symptoms of enrolled PASC patients.** Data are expressed as frequency and % of population and mean ± SEM.

**Fig. S1.**
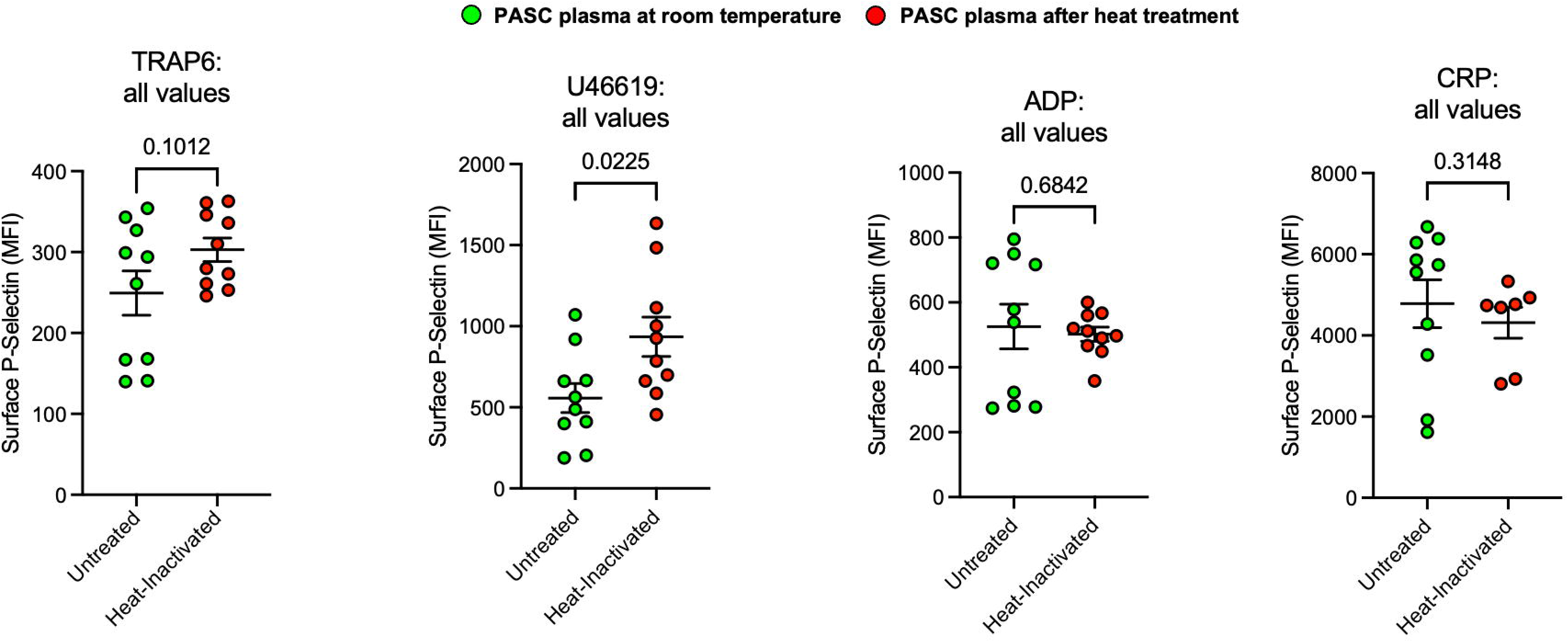
Activation of Platelets by PASC plasma persists after heat treatment. Platelet-deplete citrate plasma from PASC patients already demonstrated to activate healthy male platelets divided in half and one half was heat-treated (55°C, 15 minutes). Heat-treated and non-heat-treated plasma was incubated with washed platelets for 30 minutes prior to stimulation with a surface receptor agonist TRAP-6, 5 µM (PAR1), U46619, 1 µM (Thromboxane receptor), ADP, 0.1 µM (P2Y_12_ Receptor), or Collagen-Related Peptide, CRP, 5 µg/mL (GPVI Receptor). Platelet reactivity was then assessed by FACS by surface CD62P (P-selectin) using a tagged antibody. Data are represented as mean ± SEM. Group differences were assessed by Mann-Whitney *U* test for n=10 (room temperature citrate plasma) or n=10 (heat-inactivated citrate plasma.

**Fig. S2.**
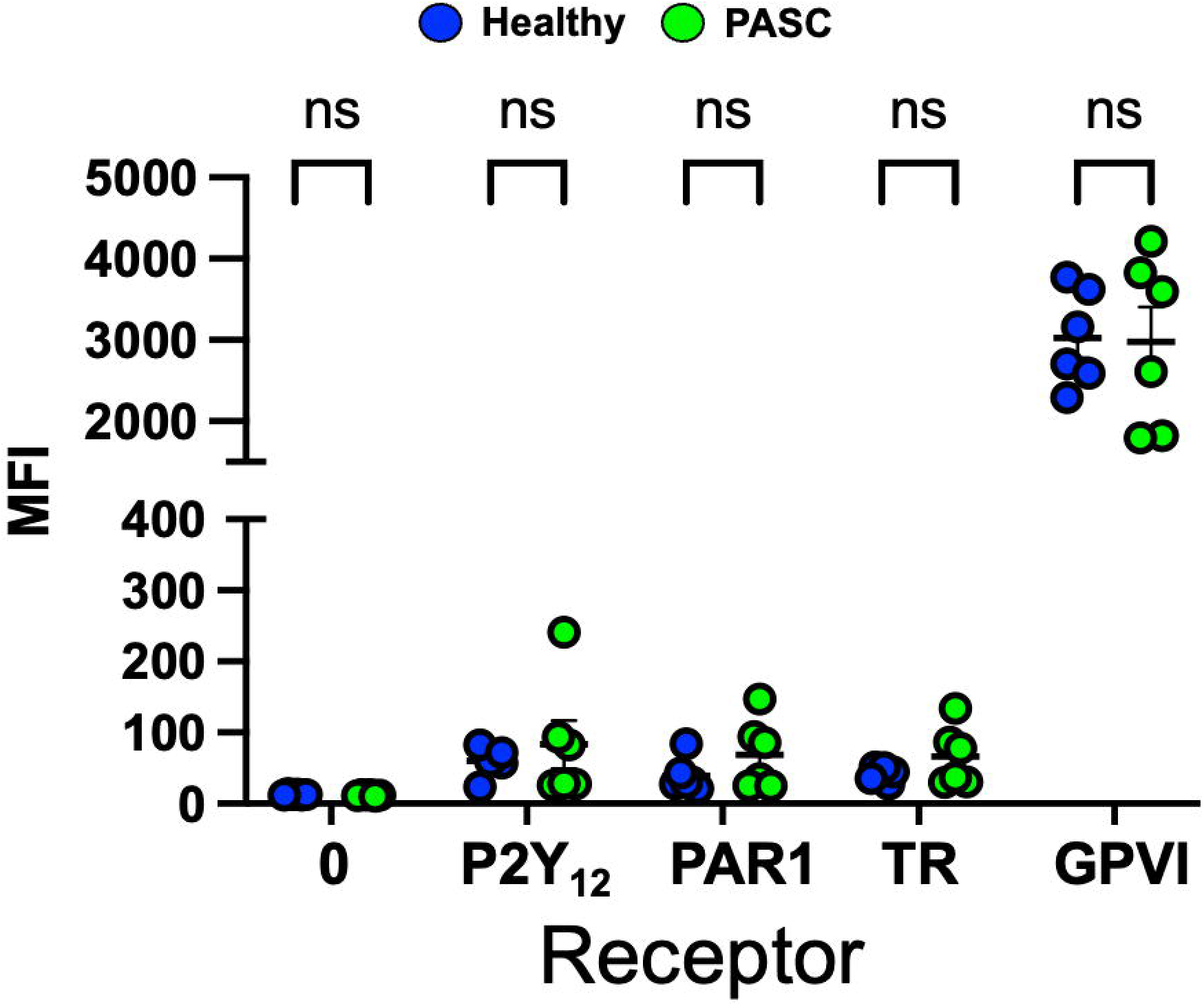
Platelet Surface Receptor Expression in PASC Patients. Blood was drawn from healthy individuals (n=6) or PASC patients (n=6) and isolated, washed platelets were assessed by FACS for expression the following receptors tagged with fluorescent antibodies: P2Y_12_, PAR1, Thromboxane Receptor (TR), GPVI, or 0 (control Immunoglobulin). Data are represented as MFI (Mean Fluorescence Intensity) of surface receptor expression ± SEM. Group differences were assessed by t-test with no differences noted between groups, ns=not significant)

**Fig. S3.**
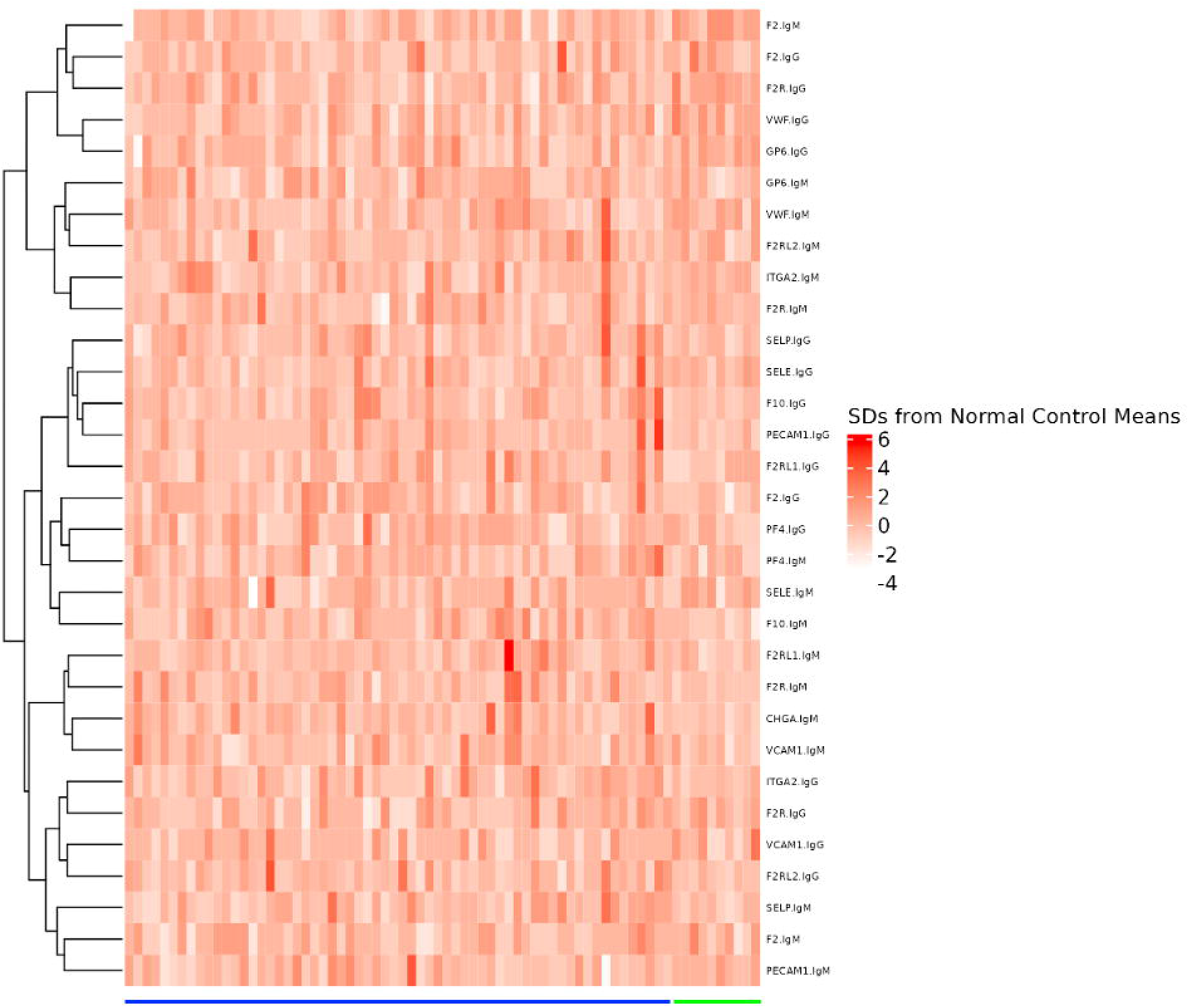
Plasma Antibody Analysis. Blood was isolated from patients with PASC (n=66, blue) or matched controls (n=10, green) and antibody profiling was performed using the HuProt 4.0 array of unique human proteins known to be exclusively expressed in platelets: Platelet Factor 4 (PF4), Glycoprotein VI (GP6), integrin alpha2 gene (ITGA2). Other proteins highly expressed in platelets were also evaluated: von Willebrand Factor (vWF), Platelet And Endothelial Cell Adhesion Molecule 1 (PECAM1), E-selectin (SELE), P-selectin (SELP), Vascular Cell Adhesion Molecule 1 (VCAM1). Coagulation cascade protein or receptors were also evaluated: Factor II (F2), factor II receptor (FIIR), Factor X (F10). A heat map was generated to show each antibody concentration expressed as standard deviations (SD) above or below the mean concentration found in healthy individuals.

**Supplemental Table 1.** Symptoms Experienced by PASC Patients.

| Symptoms | PASC (n=26) |  |
| --- | --- | --- |
|  | N | % |
| Mean Time After Infection<br>(± SEM) | 501 days ± 69 | |
| Fatigue | 20 | 76.9 % |
| Brain Fog | 20 | 76.9 % |
| Shortness of Breath | 18 | 69.2 % |
| Dyspnea on Exertion | 18 | 69.2 % |
| Dizziness | 17 | 65.3 % |
| Changes in Mood | 11 | 55.0 % |
| Insomnia | 14 | 53.8 % |
| Headaches | 13 | 50.0 % |
| Palpitations | 13 | 50.0 % |
| Chest Pain | 11 | 42.3 % |
| Joint Pain | 9 | 34.6 % |
| Cough | 7 | 26.9 % |
| Parosmia/Dysgeusia | 6 | 23.0 % |
| Memory Deficits | 6 | 23.0 % |
| Depression | 6 | 23.0 % |
| Syncope | 6 | 23.0 % |
| Nausea | 5 | 19.2 % |
| Body Aches | 4 | 15.3 % |
| Incontinence | 3 | 11.5 % |
| Neck Pain | 3 | 11.5 % |
| HF Symptoms | 2 | 7.69 % |
| Diarrhea | 1 | 3.8 % |
| Erectile Dysfunction | 1 | 3.8 % |
| Fever | 1 | 3.8 % |
| Orthopnea | 1 | 3.8 % |
| Limb Paresthesia | 1 | 3.8 % |

## Notes

### Competing Interest Statement

The authors have declared no competing interest.

### Summary of Updates

Corresponding author changed from Matthew Godwin to Scott Cameron

